# Locating highly correlated sources from MEG with (recursive) (R)DS-MUSIC

**DOI:** 10.1101/230672

**Authors:** Niko Mäkelä, Matti Stenroos, Jukka Sarvas, Risto J. Ilmoniemi

## Abstract

We introduce a source localization method of the MUltiple Signal Classification (MUSIC) family that can locate brain-signal sources robustly and reliably, irrespective of their temporal correlations. The method, double-scanning (DS) MUSIC, is based on projecting out the topographies of source candidates during topographical scanning in a way that breaks the mutual dependence of highly correlated sources, but keeps the uncorrelated sources intact. We also provide a recursive version of DS-MUSIC (RDS-MUSIC), which overcomes the peak detection problem present in the non-recursive methods. We compare DS-MUSIC and RDS-MUSIC with other localization techniques in numerous simulations with varying source configurations, correlations, and signal-to-noise ratios. DS- and RDS-MUSIC were the most robust localization methods; they had a high success rate and localization accuracy for both uncorrelated and highly correlated sources. In addition, we validated RDS-MUSIC by showing that it successfully locates bilateral synchronous activity from measured auditory-evoked MEG.

## 1 Introduction

Magnetoencephalography (MEG) offers a superior temporal resolution and a good spatial resolution to investigate the spatiotemporally complex activity of the human brain non-invasively [18]. In particular, MEG combined with source modeling offers unique information about the organization of brain functions.

Source modeling is a powerful tool for unmixing multichannel data measured with MEG or electroencephalography (EEG). To uncover brain signal sources, several methods have been developed, *e.g.*, minimum norm estimates (MNE) [16], blind source separation (BSS) techniques [4, 20], and adaptive spatial filters such as beamformers [31, 37] and multiple signal classification (MUSIC) [30, 28]. Here, we focus on the MUSIC/beamformer methods, as they exploit both temporal (via the data covariance) and spatial/physical information (via the forward model) of the measurement to solve the fundamentally ill-posed bioelectromagnetic inverse problem. As state-of-the-art magnetic resonance imaging (MRI) provides high-quality anatomical information of the human head and brain, it is reasonable to take the advantage of this anatomical information.

By “brain-signal sources”, we refer to spatially separate, macroscopically observable focal electric neuronal activity that manifests as multiple sources, and may or may not be temporally correlated between the active sites. Uncorrelated and low-to-moderately correlated brain-signal sources can be separated and located based on MEG/EEG data using multiple signal classification (MUSIC) and beamformer (BF) methods [30, 28], 37]. The conventional beamformer and MUSIC techniques, however, are limited to estimating sources that are not temporally highly correlated [2, 23]. As it is unlikely that all brain-signal sources would be uncorrelated, a special focus has been put to designing methods that can estimate sources with high temporal correlations (*e.g.,* [5, 2],[19, 9], [12, 32]). These methods, however, either require approximate prior knowledge of the locations of the sources or are designed to find synchronous, fully correlated pairs of sources. It would be useful to be able to use a single method without prior information about the source correlations.

Here, we introduce a reliable and robust MUSIC-type method for source localization, Double Scanning MUSIC (DS-MUSIC), which can locate activity consisting of a mixture of uncorrelated, moderately correlated, and synchronous, highly correlated sources. It is based on scanning over pairs of sources so that one of the sources is projected out from the data model at a time. Furthermore, we provide a recursive version of the method, RDS-MUSIC, which locates multiple sources one by one in a recursive manner, and thus, bypasses the difficult problem of peak detection present in the non-recursive MUSIC methods. This should also allow better localization of weaker sources; essentially, the recursive approach converts the difficult problem of finding multiple local maxima into a simpler problem of finding the global maximum at each consecutive recursion step [28].

A well-established way to find highly correlated sources in MEG/EEG has been presented in dual-source [2] and dual-core [9] beamformer, the latter being a technically improved version of the former. These method are designed to find pairs of synchronous sources, but their performance degrades in the presence of other non-synchronous sources. In addition, to overcome the high computational cost of the non-linear optimization scheme, the dual-source method [2] requires *a priori* information to fix the location of one of the sources, for example, with fMRI, as suggested in [2]. Dual-core beamformer (DCBF) is able to locate synchronous neural sources without this prior information [9], but the method is insufficient if some signals of interest are due to uncorrelated or weakly correlated sources.

DS-MUSIC, introduced in this study, works with any mixture of synchronous, highly correlated, correlated, or uncorrelated sources. We compared DS-MUSIC with other topographical scanning methods designed for synchronous/highly correlated sources (dual-core beamformer, DCBF, [9, 26]), or methods originally designed for uncorrelated sources that should work to some extent for highly correlated sources as well (MUSIC [30, 28], basic beamformer, BF, [37]). We performed simulations with numerous source configurations, correlation conditions, and signal-to-noise ratios (SNRs). We also compared RDS-MUSIC with the commonly used recursive method, RAP-MUSIC [28], for scanning multiple sources. Finally, we evaluated RDS-MUSIC in locating synchronous activity with measured MEG data from a well-established auditory stimulation experiment.

## 2 Theory and methods

### 2.1 Description of the measurement and scanning models

We assume that the data are generated by a finite number of brain-signal sources that can be modeled with a set of equivalent current dipoles [18]. The *measurement model* is described by

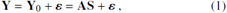

where **Y** is the measurement data matrix, **A** = [**l**_1_,…,**l***_n_*] is the mixing matrix containing the true source topographies ***l**_j_*,1,…, *n, n* is the number of sources, **Y**_0_ = **AS** the noiseless data matrix, **S** the source time-course matrix, and ***ɛ*** the noise matrix. The topographies are (*m* × 1) vectors, where *m* is the number of sensors. The noise is assumed white, *i.e.*, *εε^T^* = *σ*^2^**I**, where σ is the noise level and **I** is an identity matrix. In the case of non-white noise, we assume that the data Equation 1 can be whitened by using, *e.g.*, pre-stimulus activity or empty-room measurement [11].

We denote an equivalent current dipole by a pair (**p**, ***η***), where **p** is the location and the unit vector ***η*** the orientation of the dipole. The dipole moment is denoted with *s***η****, where *s* is the amplitude of the source. Source topographies, *i.e.*, the sensor readings due to a single source dipole are denoted by **l**_*j*_ = **l**(**P***_j_*, ***η****_j_*), *j* = 1,…, *n*, where **p**_1_,…, **p***_n_* are the source locations.

Sources are located by using a *scanning model*, which is an approximate representation of the measurement model in Equation 1. In practice, this is done by setting a discrete scanning grid over the region of interest (ROI; for example, the cortex); the scanning grid contains all source dipole candidates. A *localizer* is evaluated at all scanning grid points, and largest localizer values correspond to estimated source locations. Every dipole location **p** in the scanning grid has a (local) (*m* × 3) lead-field matrix **L(p)**, whose columns are the topographies of the source dipoles at **p** in the Cartesian *x*, *y*, and *z* directions. The measurement data are collected in the (*m* × *N*) data matrix **Y**, where *N* is the number of time points.

We assume that the columns of **A**, *i.e.*, the source topographies, are linearly independent, and thus, rank(**A**) = *n*. We also assume that the true sources have a fixed but unknown orientation. The source time-courses ***s**_i_ = **S**(i,:)* may be correlated, 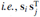 may be non-zero for *i ≠ j.* We use the Matlab notation, where **S**(*i*,:) and **S**(:, *j)* are the *i*th row and *j*th column of the matrix **S**, respectively.^1^

### 2.2 MUSIC and RAP-MUSIC

All MUSIC algorithms are based on the separation of the data space into two mutually orthogonal subspaces, the *signal space* and the *noise space*. The MUSIC scan is done by testing whether a test topography belongs to the estimated signal space or not.

Given the (unnormalized) covariance matrix of the measured data **C** = **YY**^T^, the separation to signal and noise spaces can be done by eigenvalue decomposition as follows:

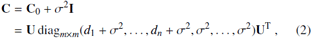

where **C**_0_ is the covariance matrix of the noiseless data, σ^2^ the noise variance, **U** contains the eigenvectors of **C**_0_, and *d_j_* are the *n* non-zero eigenvalues of **C**_0_. The approximated projection to the signal space is defined as 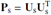, where 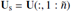 and 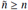 is an initial estimate for the dimension of the signal space.

In this study, we used the vector forms for the localizers, that is, the candidate source orientations were not pre-determined in the forward model [31,24]. However, all the presented localizers can be easily transformed into their scalar versions. The vector MUSIC [30] localizer is defined as

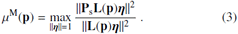

RAP-MUSIC [28] converts the MUSIC search of *n* local maxima into finding the global maximum in each successive recursion step. In every recursion, information of the previously found source estimates are projected out from the data equation with an out-projector

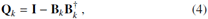

where the matrix 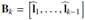 contains the topographies of the previously found source estimates, and † denotes the (Moore-Penrose) pseudoinverse. The left and right sides of the data equation are operated with the out-projector **Q***_k_*, and thus, in the next recursion, MUSIC is applied to the transformed data equation. The RAP-MUSIC localizer for the kth recursion step is given by

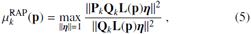

where 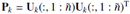 and 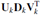 is the singularvalue decomposition (SVD) of **Q***_k_***U***_s_*. The source orientations can be determined, *e.g.*, by solving a generalized eigenvalue problem related to Eq. 5 [24].

### 2.3 Beamformers

Beamformers are a method family closely related to MUSIC [31, 29]. A conventional beamformer localizer is formed in terms of the data covariance matrix **C** and the lead-field matrix **L** that contains the candidate-source topographies of the test point.

Here, we use the conventional depth-weighted beamformer (BF), often referred to as the linearly-constrained minimum-variance (LCMV) beamformer with van Veen’s neural activity index [37, 31]. Its localizer for freely oriented sources (and assuming white noise) is given by

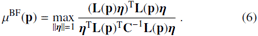

Although BF is originally derived for uncorrelated sources, it may locate also sources with relatively high correlations, given that there are no synchronous (or, linearly dependent) sources in the data. However, its performance can be decreased due to *signal loss* caused by source correlations (for more details on signal loss, see, *e.g.*, Van Veen et al. [37], Sekihara and Nagarajan [31], Mäkelä et al. [23]).

Dual-core beamformer (DCBF Diwakar et al. [9], has been suggested for locating several highly correlated and synchronous sources. It is an improved, faster version of the dual-source beamformer [2]. The DCBF localizer is given by

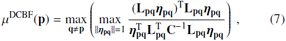

where **L_pq_** = [**L_p_,L_q_**] is the *m* × 6 dual-core topography, *i.e.*, a pair of (*m* × 3) lead-field matrices corresponding to the two sources, and ***η*_pq_** is a (6 × 1) orientation vector. Here, the DCBF localizer is written in the form of the van Veen’s neural activity index (AI) [37]. Alternatively, DCBF localizer, as well as most beamformer localizers, could be written in the pseudo-Z (PZ) format [31]. However, we chose to use the AI format, as it should be less sensitive to noise.

### 2.4 Visibility of highly correlated sources

Let us consider the data equation Eq. (1) in the case that at least two of the sources at locations **p**_1_, …, **p***_n_* are mutually highly correlated, potentially even synchronous. Sources are called synchronous if their time-courses **s**_1_, …, **s***_k_* are linearly dependent so that *α*_1_**s**_1_ + … + *α_k_***s***_k_* = 0 for *α_j_* ≠ 0; *j* = 1, …, *k*.

First, for simplicity, consider a case with n sources including a synchronous pair of sources, *i.e.*, *k* = 2. Then s_1_ = *βs*_2_ with *β= -α*_2_/*α*_1_ ≠ 0, and Eq. (1) gets the form

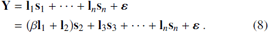

From Eq. (8) we see that the source topographies **l**_1_ and **l**_2_ do not fully belong to the signal space span(**Y**_0_) = span([*β***l**_1_ + **l**_2_, **l**_3_, …, **l**_*n*_]), and there is a combined topography 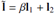 instead (by span(**M**), we denote the vector subspace spanned by the columns of a matrix **M**). We refer to this kind of nonexisting topographies as *virtual topographies*. The reasoning above implies that for the synchronous sources at **p**_1_ and **p**_2_, the localizer value may be small, *i.e.*, **p**_1_ and **p**_2_ show poorly, and can be dominated by noise. In addition, the virtual topography may lead to spurious source estimates at locations in the ROI, if some topography of the scanning model matches well with the virtual topography. Projecting out those topographies by RAP-MUSIC may remove information of the corresponding true source topographies, and consequently, they may be missed in the subsequent recursions. For example, this type of a spurious source could occur somewhere between the locations of two synchronous sources.

The reasoning above can be applied also to highly correlated sources that are not necessarily linearly dependent. As an example, let’s consider a case of *n* sources, of which two are highly correlated and located at **p**_1_ and **p**_2_ with time courses **s**_1_ and **s**_2_. Their mutual correlation coefficient 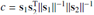 is close to one or minus one, and the time-course 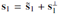, where 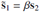 with 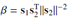, and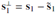 is perpendicular to **s**_2_. Now Equation 8 gets the form

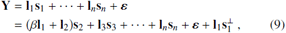

which is the equal to the Equation 8, except for the small extra term 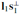, which is easily lost in noise. Then, the signal space is essentially spanned by topographies *β***l**_1_ + **l**_2_, **l**_3_,…,**l***_n_*.

To summarize, highly correlated sources show poorly, or not at all, in a MUSIC or BF scan and they may also generate spurious apparent sources. This reasoning also holds for RAPMUSIC, which will be shown in simulations. The performance of BF with correlated signal generators in MEG has been briefly discussed in [23] in the special case of two signal sources.

On the other hand, if in a recursive MUSIC method, at any step, a synchronous (or nearly synchronous) source happens to be out-projected, the mutual dependence is broken. Thus, the remaining sources are not synchronous any more and become visible and can be found. We exploit this property in developing the DS-MUSIC for locating also highly correlated sources.

### 2.5 Double scanning (DS-) MUSIC

DS-MUSIC is based on out-projecting of candidate-source topographies from the data and the scanning model in a consecutive manner so that the visibility of the correlated sources is recovered. In practice, this is done by a pairwise out-projection of a source at a *running location* **q**, while the localizer is evaluated at the *test location* **p**.

For any location **p** in the ROI, the DS-MUSIC localizer *μ*^DS^(**p**) is given by

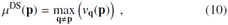

with

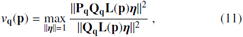

where **Q_q_** = **I**−**L(q)L(q)**^†^ is the out-projector, and the signalspace projector is 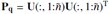, where **U** comes from **Q_q_Y** = **UDV**^T^, and 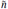 is as with the MUSIC localizer in Equation 3. Although the inner term *v*_q_(**p**) in Eq. (11) resembles the RAP-MUSIC localizer, it has two distinct differences, one in the out-projector **Q**_q_ and the other in the signal-space projector **P**_q_. First, the out-projector applies to the full (*m* × 3) topographies **L**(**q**) to project out the information related to a source at **q**, instead of the (*m* × 1) topographies in Eq. (5) with *k* ≥ 2. Therefore, it does not require the determination of the source orientations at all. Second, the signal-space projector is determined by the transformed data **Q_q_Y**, instead of the transformed signal subspace **Q_q_U**_s_ as in RAP-MUSIC. This is required to allow the breaking of linear dependence (or near dependence) of the source time-courses to take place after the transformation of the data equation. Using the **Q_q_Y** at each step is equivalent to applying the MUSIC scanning to the transformed data equation

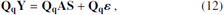

where the information of the potentially synchronous source **q** has been removed before scanning for other locations **p**.

The estimated source locations are the (largest) local maximum points of *μ*^DS^(**p**), as we see by the following reasoning. If **p** is a non-synchronous source location, then *v*_q_(**p**) ≃ 1 for each **q** ≠ **p**, and thus *μ*^DS^(**p**) ≃ 1. If **p** is a synchronous source location and **q** is the location of a synchronous partner of the dipole at **p**, then *v*_q_(**p**) ≈ 1, and thus, *μ*^DS^(**p**) ≃ 1. If **p** is not a source location, the *v*_q_(**p**) ≃ 0 for all **q** ≈ **p**, and thus *μ*^DS^(**p**) ≃ 0 (see also Table 1). Therefore, the source locations are the (largest) local maximum points of *μ*^DS^(**p**). The computational load of DS-MUSIC is of order *K^2^* where *K* is the number of scanning grid points.

**Table 1.** Working principle of DS-MUSIC localizer.

| Test point $\mathbf{p}$ | Correlation | Running point $\mathbf{q}$ | $\nu_{\mathbf{q}}(\mathbf{p})$ | $\mu^{\text{DS}}(\mathbf{p})$ |
| --- | --- | --- | --- | --- |
| Source point | Synchronous | Not a pair | $\approx 0$ | $\approx 0$ |
| Source point | Synchronous | Is a pair | $\approx 1$ | $\approx 1$ |
| Source point | Uncorrelated | - | $\approx 1$ for all $\mathbf{q}$ | $\approx 1$ |
| Not source point | - | - | $\approx 0$ for all $\mathbf{q}$ | $\approx 0$ |

### 2.6 Recursive DS-MUSIC (RDS-MUSIC)

Here we show how to expand DS-MUSIC into a recursive approach that we call RDS-MUSIC. This is done in order to gain similar benefits as RAP-MUSIC has in comparison to MUSIC. In other words, RDS-MUSIC searches one global maximum at each recursive step instead of multiple local maxima in one DS-MUSIC run.

The first recursion is equivalent to the non-recursive DSMUSIC scan; the first source-location estimate **p**_*i*_1__ is obtained as the global maximum point of the localizer *μ*^DS^(**p**). After locations **p**_*i*_1__,…, **p**_*i*_*k*-1__ have been found, one forms the matrix **B**_*k*_ = [**L**(**p**_*i*_1__),…,**L**(**p**_*i*_*k*-1__)] and with that, the localizer 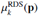 for the *k*th iteration step by

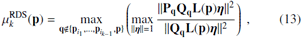

for 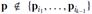, Where 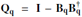 with **B**_q_ = [**B**_k_,**L**(**q**)], and 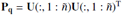, Where **U** is obtained from **UDV**^T^ = **Q_q_Y**, and 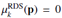 for the already-found locations **p** ∊ {**p**_*i*_1__,…, **p**_*i*_*k*-1__}.

The iteration is continued until all source dipoles have been found, which is typically indicated by a significant drop of the maximum value of 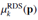, or *post hoc*, after 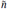 recursions, by using the drop of the maximum value of 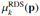 as a classifier for true *vs*. false sources [28, 24]. The classification can be done either by setting a fixed threshold for the localizer value maximum [28, 22]), or adaptively, by monitoring the evolution of the localizer value maximum as a function of recursion steps. In general, the adaptive approach should be more robust [21, 3, 24].

The computational cost of RDS-MUSIC is *nK^2^* (or, in practice, 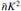 with 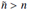), where *n* is the number of sources and *K* the number of scanning grid points; thus, the computation time of RDS-MUSIC scales linearly with the number of sources.

## 3 Simulations and measurements

In this section, we describe the simulations that provide *(i)* illustrative example comparisons of our new methods with other source localization methods (MUSIC, RAP-MUSIC, and beamformers) in Simulations 1 and 3, and *(ii)* statistical evidence from extensive comparisons of different methods in Simulations 2 and 4. The simulations and analysis of both simulated and measured data were carried out with Matlab (The Mathworks Inc., Natick, MA, USA). We compared non-recursive methods (MUSIC, DS-MUSIC, BF, and DCBF) and recursive methods (RDS-MUSIC and RAP-MUSIC).

### 3.1 The head models

The simulations were done in a realistic head geometry, using different head models to compute the lead fields for the “true” and “scanning” models.

A scanning grid with a spacing of 4 mm and 871 grid points was set to an axial plane inside the cranium, 4 cm below the top of the head (see 1). Similar approach has been used previously in [24] and [27]. The MEG forward solution for the source candidates in the scanning grid was computed by a linear-collocation boundary-element method (BEM) with the isolated-source approach in a 3-layer head model including boundary surfaces for the scalp, skull, and brain [17,34],35]. The conductivity values for the scalp, skull, and brain surfaces were 0.33 (Ωm)^-1^, 0.0066 (Ωm)^-1^, and 0.33 (Ωm)^-1^, respectively [14, 6]. For the statistical comparisons (Simulation 2 and 4), we used a sparser scanning grid with an 8-mm spacing and 221 grid points.

**Figure 1.**
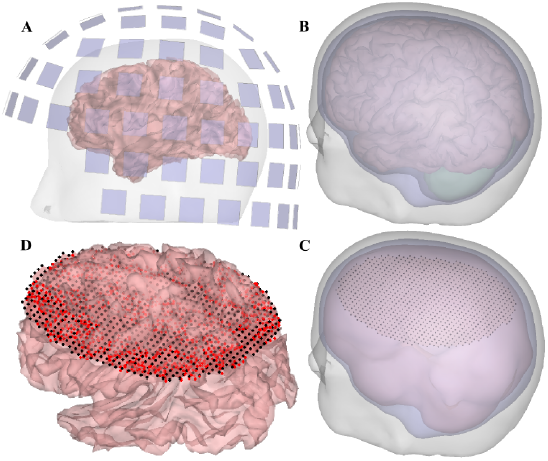
The head-model geometries. (**A**) shows the MEG sensors, scalp, and cortex, (**B**) the ‘true’ 4-compartment model, (**C**) the ‘ scanning ’ 3-shell BEM models accompanied with the scanning grid (black dots), and (**D**) shows all possible locations of the true (red dots) sources, and the scanning grid points (black dots) visualized on the cortical surface.

The true sources were sampled from a triangular mesh with a spacing of 3.5 mm. The mesh was overlaid on the cortical surface, and a dipole was set at each point. The dipoles were perpendicular to the cortical surface, pointing outwards from it. The topographies for these sources were computed with a linear collocation with the isolated-source approach in a 4-compartment boundary-element model [34, 33] with compartments for the scalp, skull, cerebrospinal fluid (CSF), cerebellum, and cerebral cortex (see 1). Commonly used conductivity values for the scalp, skull, CSF, and brain were used: 0.33 (Ωm)^-1^,0.0066 (Ωm)^-1^, 1.79 (Ωm)^-1^, and 0.33 (Ωm)^-1^, respectively [1, 14], 33]. Only those points of the mesh that were closer than 3 mm to the scanning grid were selected as the final simulation space, yielding a subset of 1075 dipole locations, from where the true sources were selected for each simulated data set. The true sources were drawn randomly from these ROI points so that they were at least 3 cm apart from each other. This selection was done to ensure that the sources were initially spatially separable because the scope of this study was to investigate the effect of their *temporal* similarity on their visibility in different MUSIC and beamformer scans. The 4-compartment model was created as in [33], using a sample data set of SimNIBS [40]. There were 102 sensor units in the simulated MEG system; each unit had two planar gradiometers and one magnetometer, as in the 306-channel MEG of Elekta Neuromag. White noise was added to the noiseless data according to Eq. (1) in all simulations. The magnetometer noise was scaled with a factor of ten so that all sensors had approximately the same noise level.

### 3.2 Simulation 1: Comparison of DS-MUSIC with MUSIC, BF, and DCBF in two example data sets

Here we demonstrated the performance of our DS-MUSIC in two simulated data sets that contained both synchronous and uncorrelated sources, and compared DS-MUSIC with the conventional MUSIC [30, 28] and BF [37], as well as with DCBF [9].

The data sets had a synchronous pair of sources, and one or two uncorrelated sources (yielding *n* = 3 and *n* = 4, respectively). All time-courses were sinusoidal. Note, however, that the form of the time-courses is actually not essential, because in the input of MUSIC and BF methods only the source covariance matrix **C**_s_ = **SS**^T^ matters. We selected 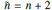 for all MUSIC-type methods.

### 3.3 Simulation 2: Statistical comparison of DS-MUSIC with MUSIC, BF, and DCBF

We compared DS-MUSIC with non-recursive MUSIC, BF, and DCBF in numerous situations with varying key parameters. We hypothesized that MUSIC and BF should perform well with data sets due to only uncorrelated/moderately correlated sources, DCBF with only synchronous sources, and DS-MUSIC in all situations.

To compute the quality measures for localization accuracy of the non-recursive localizers, the source location corresponding to the global maximum of the concerned localizer was identified, and then, its neighborhood with a radius of 3 cm was ignored in finding the next global maximum. This procedure was repeated to obtain *n* peaks, where *n* was the number of simulated sources. All sources were considered successfully located, if each source estimate was located closer than one cm from the corresponding true source. Each method could thus gain one or zero points from each simulated data set, so that the maximum success rate (SR) for 100 simulations was 100. For a single data set, mean localization error of the estimated *n* sources was computed, and these values were averaged over 100 simulations to obtain the overall localization error (LE).

The performance of the methods was compared in 8 different source configuration types (SCTs) with different mutual correlations between the sources. The studied SCTs were

- for *n* = 3: uncorrelated sources, weakly correlated sources, highly correlated sources, a synchronous pair + one uncorrelated source, and a synchronous triplet + one uncorrelated source,
- for *n* = 4: a synchronous pair + two uncorrelated sources, and a synchronous triplet + one uncorrelated source, and
- for *n* = 2: a synchronous pair.

With a synchronous pair, we refer to a situation with, *e.g.*, **s**1 = *α***s**_2_, and with a synchronous triplet, to **s**_1_ = *α***s**_2_ +*β***s**_3_, with **α*,*β* ≠* 0. We did not consider situations with three (or more) sources with exactly parallel time-course vectors, *i.e.*, **s**_1_ = *α***s**_2_ = **β***s**_3_*, *α*,*β* ≠ 0. Such a situation would most likely cause problems to all of the studied methods, but is probably unlikely in real-life MEG/EEG measurements. Instead, we considered a more realistic case with three sources with identical sinusoidal time-courses with small mutual phase shifts, similarly as in [9] (see Simulations 3 and 4 for details).

Weak correlation was set to 0.1–0.3, and high correlation to 0.5–0.9. A synchronous pair had a time-course **s**_1_ = **s**_2_ and a synchronous triplet **s**_1_ = **s**_2_ + **s**_3_. All source time-courses were first simulated as sinusoids, or as a sum of sinusoids for the triplet; thereafter, their mutual correlations were set to a desired level by multiplying with a preset correlation matrix or by creating phase shifts between the time-courses. All simulations were repeated with SNRs of 2, 1, and 0.75. The SNR was defined as the ratio of the Frobenius norms of the noiseless part of the signal and the noise matrix. We used 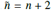 2 for all MUSIC-type methods.

### 3.4 Simulation 3: Comparison of RDS-MUSIC vs. RAP-MUSIC in three example data sets

Here we demonstrated the performance of RDS-MUSIC and compared it with the well-established recursive source localization method, RAP-MUSIC [28], in three example data sets with different source configurations. SNR was set to one for all cases and 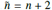 for both methods.

The three simulated test cases were

- Case 1: *n* = 2 with a synchronous pair of sources. This example demonstrates the simplest case with synchronous sources.
- Case 2: *n* = 3 with a synchronous pair and one uncorrelated source. This example demonstrates the simplest case with both synchronous and uncorrelated sources.
- Case 3: *n* = 3 with three sinusoidal time-courses with identical frequency but different phase angles of 0,5°, and 10°, so that the time-courses are very highly correlated (correlation coefficient > 0.98).

### 3.5 Simulation 4: Statistical comparison of RDS-MUSIC vs. RAP-MUSIC

Here, we computed statistical measures from numerous simulations comparing RDS-MUSIC and RAP-MUSIC. We analyzed 100 simulated measurements for each of the five source configurations: *(i) n* = 2 with a synchronous source pair, *(ii) n* = 3 with uncorrelated sources, *(iii)* with phase-shifted signals of the same frequency, *(iv)* with a synchronous pair + one uncorrelated source, and *(v) n* = 4 with a synchronous pair + two uncorrelated sources.

SR and LE were used as quality measures for both methods, as in Simulation 2. Now, contrary to Simulation 2, no peak detection was necessary. Instead, the estimated source locations were determined directly by the scanning grid point that corresponded to the global maximum of the localizer at each recursion step. The differences of the localization errors were compared with the Mann-Whitney U-test with an error rate of 0.001. The statistics were adjusted for multiple comparisons with Bonferroni correction.

### 3.6 Measured MEG data

We recorded MEG data during auditory stimulation of a 29-year-old right-handed neurologically normal man. The subject gave a written informed consent. The experiments were performed in accordance with the Declaration of Helsinki, and they were permitted by the Research Ethics Committee of Aalto University. The measurements were done with a 306-channel MEG scanner (Elekta Neuromag, Elekta Ltd., Helsinki, Finland), located in the magnetically shielded room (IMEDCO AG, Hagendorf, Switzerland) in Aalto Neuroimaging MEG Core.

Auditory stimulation was performed by delivering short pure tones monoaurally to the left or right ear. These stimuli are well-established and known to produce synchronous activity in left and right auditory cortices (*e.g.*, Virtanen et al. [38]). The volume of the sinusoidal audio input was set to a comfortable level before the measurement.

The data were collected with a sampling frequency of 1000 hertz. 160+160 epochs were recorded from a randomized trial-order with an inter-stimulus interval of 2.6–3 s. Signal-space separation [36] and interpolation of channels were applied to the data with the Maxfilter software (Elekta Ltd.). The data were band-pass-filtered to 2–80 Hz and a quality check of channels and epochs was carried out. Data were averaged over the epochs. No whitening was applied, as MUSIC has been shown to work well without whitening for measured event-related MEG [24]. A time window of 10-110 ms with respect to the stimulus triggers (at time *t* = 0) was analyzed.

The individual head model was computed from the subject’s T1-weighted MRIs using Freesurfer and MNE software [13, 15]. Three layers of boundary meshes were obtained. A boundary-element model (BEM) was constructed using commonly used conductivities of 0.33 (Ωm)^−1^ for the scalp and brain and 0.0066 (Ωm)^−1^ for the skull were used [35]. The cortical source space had 10242 grid points per hemisphere with approximately 3.1 mm spacing. The forward model without *a priori* information about source orientations was computed with the BEM based on linear Galerkin boundary elements, formulated with the isolated-source approach [17, 34].

## 4 Results

### 4.1 Simulation 1: Examples with DS-MUSIC

Two case examples with a synchronous pair plus one uncorrelated source and a synchronous pair plus two uncorrelated sources are visualized in fig2. These cases illustrate what typically happened in our simulations with a synchronous source pair accompanied with one or two other sources. fig2 with the localizer-value distributions and their local maxima visualized on the ROI shows that in cases with mixtures of uncorrelated and synchronous sources, only DS-MUSIC was able to locate all sources. MUSIC, BF, and DCBF accurately located only a subset of the sources. The results of this simulation benchmark the previously derived theory and verifies our implementations of the methods.

**Figure 2.**
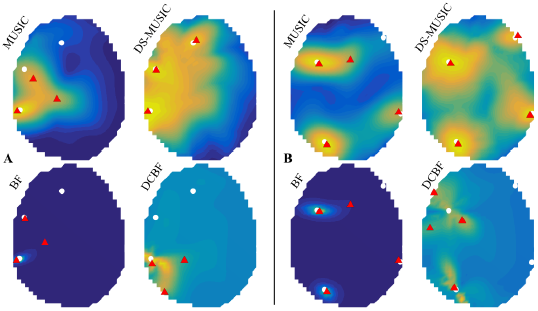
Two example cases with (**A**) *n* = 3 and (**B**) *n* = 4, both including a synchronous source pair. The scanning results of MUSIC, DS-MUSIC, BF, and DCBF are visualized with a colormap (yellow color corresponds to high localizer values). The true source locations are indicated with white dots, and the *n* localizer maxima peaks with red triangles.

### 4.2 Simulation 2: Statistical results with DS-MUSIC

The statistical results of the comparisons of MUSIC, BF, DCBF, and DS-MUSIC are shown in fig3. MUSIC and BF performed well with high success rates (SR) and low localization errors (LEs) in cases with uncorrelated and weakly correlated sources. They also performed relatively well in the case of sources with mutual correlations between 0.5–0.9. MUSIC and BF did not tolerate synchronous sources. DCBF performed well only in the cases with a synchronous pair or a synchronous triplet. DS-MUSIC had high success rates in all simulation situations; it gave high SRs and LEs independent of the correlation levels of the sources.

**Figure 3.**
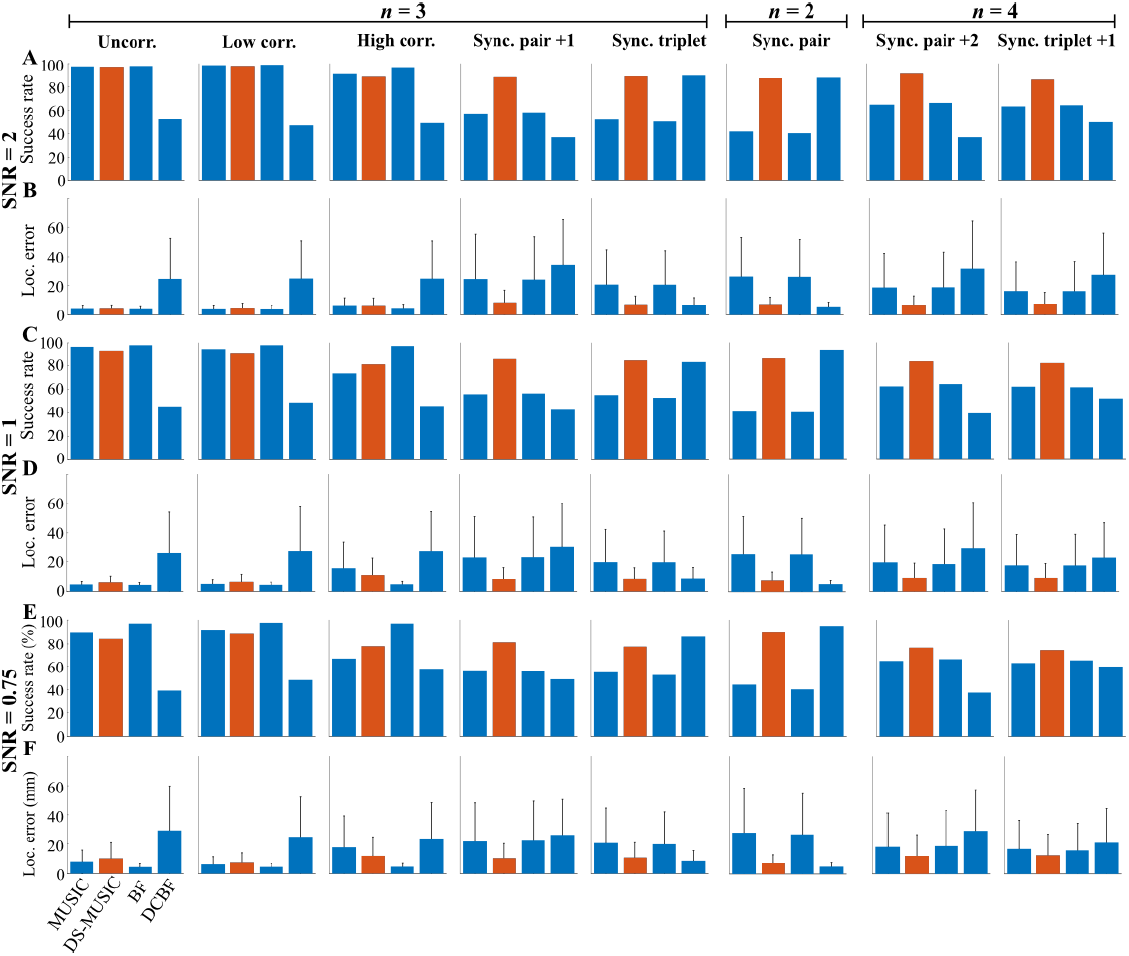
Success rates and localization errors from comparisons of MUSIC, DS-MUSIC, BF, and DCBF, with 3 different SNRs and 8 different source configuration types. 100 cases were simulated for each condition. DS-MUSIC is visualized in orange and other methods with blue. Success rates and mean localization errors are in (**A,C,E**) and (**B,D,F**), respectively. Standard deviations of the localization errors are indicated with black lines.

### 4.3 Simulation 3: Examples with RDS-MUSIC

We analyzed three different case types. Example cases, representing a typical result in each case type, are shown in fig4–fig6.

Case 1 had *n* = 2 with a synchronous pair of sources. The localization results are shown in fig4. RDS-MUSIC was able to locate both sources, whereas RAP-MUSIC successfully found only one source. RAP-MUSIC estimated a false source located between of the two synchronous sources, as predicted by the theory.

**Figure 4.**
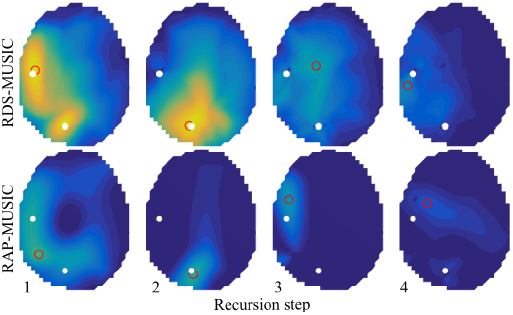
Localization results of Case 1 with *n* = 2 with a synchronous source pair. The localizer value distribution is visualized on the ROI with a colormap. True sources are marked with white circles and their estimates with red rings.

Case 2 with *n* = 3 had a synchronous pair and one uncorrelated source (fig5). RDS-MUSIC successfully located all 3 sources, and gave a strong contrast between the true (*i.e.*, 3 first recursions) and false sources (*i.e.*, 2 last recursions). RAP-MUSIC located only the uncorrelated source accurately, and suggested a spurious source in between of the two synchronous sources.

**Figure 5.**
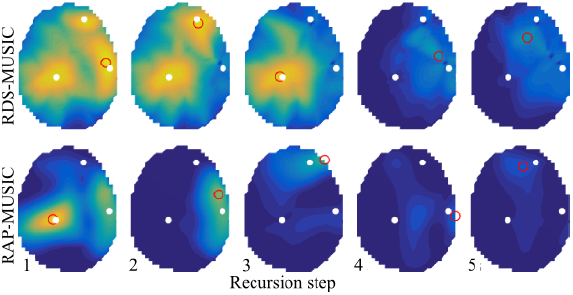
Localization results of Case 2 with *n* = 3 with a synchronous source pair. The localizer value distribution is visualized on the ROI with a colormap. True sources are marked with white circles and their estimates with red rings.

Case 3 with *n* = 3 sources with matching frequency but different phase angles demonstrated the performance of RDS- and RAP-MUSIC on highly correlated but not synchronous sources (6). RDS-MUSIC successfully located all three sources with very good accuracy, and did not suggest any spurious sources. RAP-MUSIC located only two out of three sources successfully. The accuracy of RAP-MUSIC diminished after the first source.

**Figure 6.**
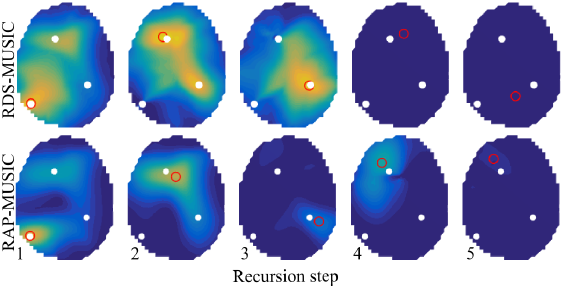
Localization results of Case 3 with *n* = 3 with three highly correlated sources. The localizer value distribution is visualized on the ROI with a colormap. True sources are marked with white circles and their estimates with red rings.

The maximum value of the localizer as function of recursion steps is shown in 7 for Cases 1-3. This value is typically used in estimating the number of sources in RAP-MUSIC [28, 24]. In all cases, RDS-MUSIC gave a good contrast between the true (2 or 3, depending on the case) and false sources (*i.e.*, the additional 2 recursion). RAP-MUSIC did not show good performance in estimating the number of sources, and failed with Case 3 with phase-shifted source signals (7C).

**Figure 7.**
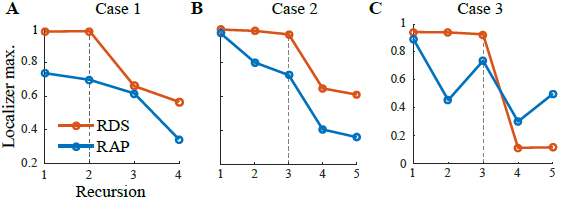
Estimation of the number of sources. Maximum values of the RDSMUSIC (orange) and RAP-MUSIC (blue) localizers as functions of recursion steps are from Cases 1–3 (see Figs. 4–6). The true n is indicated with a vertical dashed line.

### 4.4 Simulation 4: Statistical results with RDS-MUSIC

The SR and LE comparisons of RDS-MUSIC and RAP-MUSIC are shown in 8. The results show that with uncorrelated sources, the methods performed equally well, as predicted by the theory. However, with synchronous sources, potentially accompanied with one or more uncorrelated sources, RDS-MUSIC outperformed RAP-MUSIC in all scenarios and SNRs. With the case of three sources with phase-shifted sinusoidal time-courses, RDS-MUSIC was significantly better than RAP-MUSIC with SNRs of 1 and 0.75.

**Figure 8.**
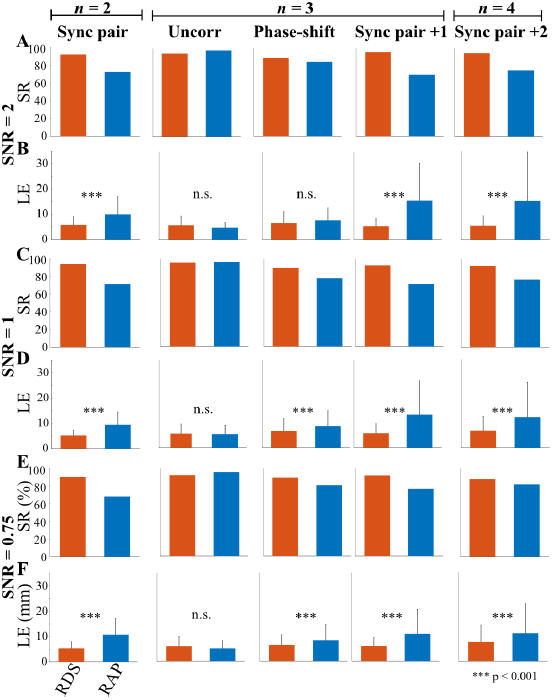
ESuccess rates and localization errors from comparisons of RDSMUSIC and RAP-MUSIC, with 3 different SNRs and 5 different source configuration types. DS-MUSIC is visualized in orange and RAP-MUSIC with blue. Success rates and mean localization errors are in (**A,C,E**) and (**B,D,F**), respectively. Standard deviations of the localization errors are indicated with black lines.

### 4.5 Measured MEG data

Auditory evoked MEG data were analyzed with RDS-MUSIC and RAP-MUSIC (9). Results are shown in 9. RDS-MUSIC was able to locate accurately two sources bilaterally, one in the left and one in the right primary auditory cortex. RAP-MUSIC was able to locate only one source correctly; it was located close to the left auditory cortex. RAP-MUSIC located the second source close to the geometric origin of the cortex model. The SNR of the measured MEG was 4, determined as the ratio of the Frobenius norms of the data and noise matrices. The noise matrix was collected from a time window of −100–0 ms prestimulus.

**Figure 9.**
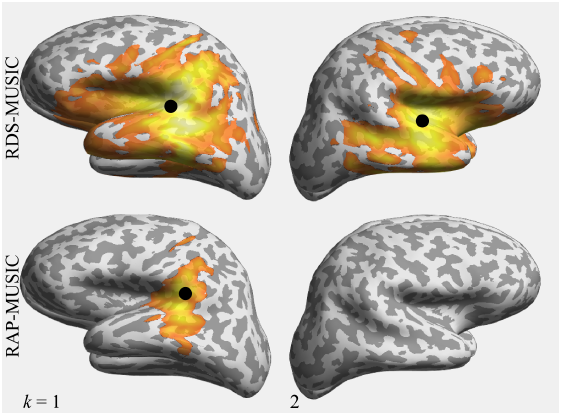
Localization results for the measured auditory-evoked MEG, analyzed with RDS-MUSIC and RAP-MUSIC. RDS-MUSIC located two sources bilaterally accurately to the auditory cortices. RAP-MUSIC located only one source, close to the left auditory cortex. The localizer values above 0.75 are visualized with the ‘ hot ’ colormap, and the maximum values of the localizers, *i.e.*, the estimated source locations, are marked with black circles on the inflated cortical surface of the subject.

## 5 Discussion

We introduced the DS-MUSIC algorithm for locating sources regardless of the level of correlation between the source time-courses. In addition, we provided its recursive version, RDS-MUSIC, to bypass the challenging multiple-local-maxima search present in the non-recursive algorithms. Our localization methods performed with high success rates in simulations with various source configurations, temporal correlations of the sources, and within a realistic range of SNRs. In addition, RDS-MUSIC successfully located a bilateral primary auditory cortex activity with good accuracy from real MEG during sound stimulation.

Therefore, we argue that DS-MUSIC and RDS-MUSIC are robust and powerful means for investigating brain activity, where some signal generators may be temporally correlated while some are not. The simulations and measurements were done for MEG, but in principle, the proposed methods should work with EEG as well.

We compared DS-MUSIC with other non-recursive source localization techniques; MUSIC, BF, and DCBF. DS-MUSIC worked well with *(i)* zero to moderately correlated sources, with similar performance as the conventional MUSIC and BF, *(ii)* highly correlated and synchronous sources, *(iii)* as well with mixtures of *(i)* and *(ii*). On the other hand, the performance of MUSIC and BF deteriorated with data due to highly correlated sources, especially with lower SNRs, and they failed when synchronous sources were present. These results validated our theoretical reasoning in Theory and methods: the visibility of the sources is decreased when they are highly correlated, and the small visible parts of the signals can be lost in noise. The DCBF algorithm, on the other hand, performed well only when a pair or triplet of synchronous sources were present, and failed if the data were also due to some uncorrelated sources. To conclude, DS-MUSIC provided the most robust performance over different correlation conditions of the sources.

Non-recursive techniques such as MUSIC and BF suffer from being suboptimal for multiple source localization [28]—this applies also to DS-MUSIC. Therefore, we extended our DS-MUSIC in a recursive approach, RDS-MUSIC. The recursive method provides similar benefits as the well-established RAP-MUSIC compared to MUSIC [28], but the DS-approach additionally allows better performance when highly correlated and synchronous sources are present (Simulation 3; Figs. 4–7, and Simulation 4; 8). RDS-MUSIC outperformed RAP-MUSIC in analyzing MEG due to synchronous auditory-evoked activity; RDS-MUSIC located two sources accurately in the primary auditory cortices, whereas RAP-MUSIC located only the other, and did that with a relatively poor accuracy (9).

Although the assumption of uncorrelated sources is not explicitly used when deriving the basic MUSIC algorithm—contrary to basic beamformers—high correlations clearly degraded the performance of MUSIC and RAP-MUSIC as well. Indeed, the methods tolerated correlations to some level, but they did not perform well with data with synchronous sources. This result is supported by earlier studies, where RAP-MUSIC has been shown to produce spurious sources between of two synchronous sources [25], just like conventional beamformers [23].

As some methods are designed for uncorrelated sources and some for highly correlated sources, one could argue that applying a two-step process of *first* locating uncorrelated sources with a suitable technique and *then*, locating correlated sources with another suitable technique (or *vice versa*), could be used for a mixture of sources with varying correlations. Our results suggest, however, that this would not be a good choice. This can be seen from Figs. 4–6; a synchronous source pair can destroy the localization of uncorrelated sources, and an uncorrelated source can destroy the source localization of synchronous sources, with the result that spurious “sources” are generated. Therefore, when one cannot be sure of the correlation conditions of the signal generators in MEG (or EEG), we would recommend to use (R)DS-MUSIC for scanning multiple sources.

We recently presented a modified version of RAP-MUSIC [24]. Its improvements were specifically targeted to make the estimation of the number of uncorrelated or weakly correlated sources more accurate. Therefore, we did not include that method in this study, but decided to concentrate on well-established methods for comparison. Based on the theoretical foundation of TRAP-MUSIC, source correlations should affect its behavior in the same way as for RAP-MUSIC, and hence, RDS-MUSIC is necessary if data is partly due to highly correlated sources. We also did not include the so-called enhanced DCBF [10] to this study, as we did not observe any significant differences between DCBF and eDCBF in our test simulations that were similar to Simulation 2 with SNR of two.

DS-MUSIC and RDS-MUSIC exploit the 3-D local lead-field matrix **L**(**p**) instead of the 1-D source topography vector in the out-projections. Therefore, the methods should be less sensitive to the errors in segmenting and discretizing the cortex, and thus, the orientation of the candidate sources. When a candidate source at a location **p** has a pre-determined orientation that does not match well with the true orientation at or nearby that location, the localizer value gives smaller values than it should. On the other hand, using the full **L**(**p**) decreases the spatial resolution of the method slightly, as it projects out a larger subspace. If one is interested in the source orientations, they can be estimated subsequently with, *e.g.*, the approach presented by Shahbazi et al. [32].

We simulated the true source topographies with a state-of-the-art 4-compartment model that has a realistic CSF compartment, and estimated the sources using a typical 3-shell model, where the CSF is omitted. In addition, the grids for true sources and scanning space were chosen different. As inaccuracies are always present in any real-life MEG/EEG source-level analysis, these choices bring realism to our simulations. If these modeling errors were omitted, it could lead to unrealistically optimistic results or introduce bias favoring some method over the other ones.

The computational efficiency of the DS-MUSIC is dominated by the size of the scanning grid, which defines the number of candidate topographies. Thus, the computational load grows rapidly with denser scanning grids, making the method relatively slow. The computational load of DS-MUSIC is of the same order as that of DCBF [9], which constructs dual-core topographies, *i.e.*, pairs of candidate topographies, for the scanning. These methods can be, however, made faster if necessary. For example, sparser scanning grids could be used. Also other techniques could be used to obtain faster computation times. For example, conjugate-gradient or Powell’s search [9], forward models with pre-determined fixed source orientations [39, 31], or smart topographical clustering [7, 8], could be applied in conjunction with the actual source localization method. A set of only 100–200 topographies have been shown relatively sufficient to describe the whole cortex, when combined with an atlas-based clustering (see, *e.g.*, Dinh et al. [8]). This approach seems reasonable, in particular in online analysis, as the spatial resolution of MEG (or EEG) is anyway of the order of several millimeters at best.

## 6 Conclusion

We introduced the DS-MUSIC method, which is suitable for locating sources regardless of their potential temporal correlations. We compared it with MUSIC, beamformer and DCBF, showing that it was the most robust localizer across different source configurations. We expanded DS-MUSIC to a recursive RDS-MUSIC and compared it with RAP-MUSIC. As predicted by theory, RDS-MUSIC performed better than RAP-MUSIC when signals of interest were partially or completely due to synchronous sources or highly correlated sources, and was as successful with uncorrelated sources as RAP-MUSIC. When applied to measured auditory evoked MEG, RDS-MUSIC located sources bilaterally to the primary auditory cortices. Thus, (R)DS-MUSIC is a robust and powerful tool for locating brain activity, especially when there is a good chance that at least some of the signal sources are highly correlated.

## Acknowledgments

This study was funded by Academy of Finland (grant number 283105), Foundation for Aalto University Science and Technology, Orion Research Foundation, and KAUTE Foundation.

## Footnotes

1 1We have used similar notation and description of the data (when applicable) as in [24].

